# Sleep regulation of the distribution of cortical firing rates

**DOI:** 10.1101/084731

**Authors:** Daniel Levenstein, Brendon O. Watson, John Rinzel, György Buzsáki

## Abstract

Sleep is thought to mediate mnemonic and homeostatic functions. However, the mechanism by which this brain state can implement both the “selective” plasticity needed to consolidate novel memory traces as well as the “general” plasticity necessary to maintain a well-functioning neuronal system is unclear. Recent findings show that both of these functions differentially affect neurons based on their intrinsic firing rate, a ubiquitous neuronal heterogeneity. Furthermore, they are both implemented by the NREM slow oscillation, which also distinguishes neurons based on firing rate during sequential activity at the DOWN->UP transition. These findings suggest a mechanism by which spiking activity during the slow oscillation acts to maintain network statistics that promote a skewed distribution of neuronal firing rates, and “perturbation” of that activity by hippocampal replay acts to integrate new memory traces into the existing cortical network.

## Introduction

Extensive work on non-REM (NREM) sleep has revealed two major functions: memory consolidation [1] and homeostasis [2,3]. These functions are each thought to result from synaptic plasticity, but are directed towards two distinct goals. In the case of memory consolidation, specific synapses are thought to be “selectively” modified to strengthen particular memory traces. On the other hand, homeostatic function is thought to involve a “general” population-wide modification of synapses to maintain a stable neuronal system. While these plasticity mechanisms are often considered independently, recent findings indicate that both homeostasis and memory consolidation exhibit differential effects on neurons with respect to a ubiquitous neuronal heterogeneity-intrinsic firing rate. During homeostatic conditions, higher firing rate excitatory cells were found to exhibit a decrease in spiking activity while those with lower firing rates increase their spike rate [4,5]. On the other hand, after a learning task induced memory consolidation, recordings showed replay of spike sequences in hippocampal pyramidal cells wherein the tuning of higher firing rate cells is “rigid”, or relatively unchanging with behavioral experience, while low firing rate neurons act as a pool of “plastic” cells available to be incorporated into novel memory traces [6]. These findings are of particular interest given that spontaneous firing rates in these populations are distributed over three orders of magnitude in a highly skewed distribution, suggestive of a systematic overarching organization that allows for structured functional diversity *within* a local population of neurons of the same cell type [7] (Figure 1A).

**Figure 1.**
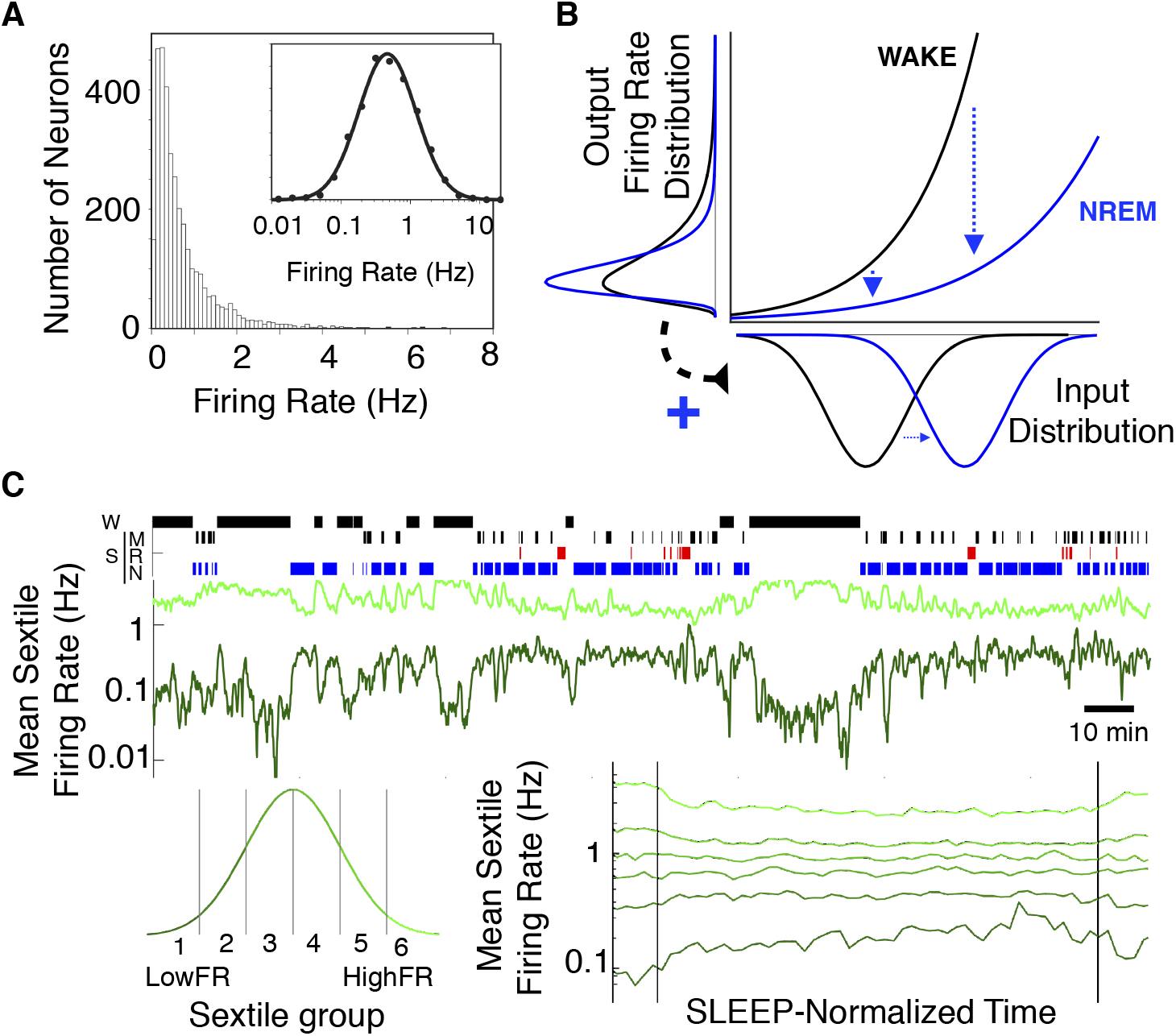
Skewed distribution of firing rates in sleep and wake. **A**: Skewed distribution of spontaneous firing rates on a linear and log scale. (from Mizuseki and Buzsáki 2013) **B**: Supra-linear neuronal input-output (F-I) curve as source of the firing rate skew and mechanism for within-NREM narrowing of the firing rate distribution. Loss of neuromodulatory tone during NREM (i) decreases the gain of the F-I curve and (ii) increases the strength of local connections. Together, the expected result is to narrow the distribution of firing rates. (adapted from Roxin et al 2011). **C**: Firing rate of high and low rate sextile groups over wake (W) and sleep (S) states (M: Microarousals, R: REM, N: NREM) for an example recording session (top), and mean of all firing rate sextiles during sleep over all recordings (bottom).

In this review, we discuss how the neocortical NREM slow oscillation can mediate plasticity towards both homeostatic and mnemonic goals. We first review how sleep state differentially affects neurons as a function of the biophysical heterogeneities that result in firing rate variability across the population. Based on the application of synaptic plasticity rules to observed NREM population dynamics, we then propose a mechanism by which NREM sleep enacts homeostatic maintenance of the neuronal firing rate distribution. This novel homeostatic mechanism is necessary to counteract a widening distribution of firing rates, which is a consequence of wake-like regimes. Together, these effects maintain general network statistics that have beneficial properties for mnemonic function and network stability. Lastly, we discuss how perturbation of the activity pattern responsible for this general homeostatic modification by hippocampal replay events can result in the selective modifications necessary for memory consolidation.

## Biophysical heterogeneity as a source of the neuronal firing rate distribution

Heterogeneity, or variability in neuronal properties across a population, is a biological reality that has profound implications for the dynamics and function of neural circuits [8]. In one ubiquitous example of heterogeneity, mean spike rates of individual neurons are distributed over three orders of magnitude in a highly skewed lognormal distribution [7](Figure 1A). This distribution in firing rates is seen during both “spontaneous” and “evoked” activity [9,10], and individual neurons maintain their rank in the firing rate over the course of months and across behavioral states [9,11]. Furthermore, a given neuron will return to a firing rate set-point in response to long-term perturbation of input [12]. Together, these observations suggest that the distribution of intrinsic firing rates reflects a fundamental *biophysical* heterogeneity in neuronal populations, and that the mean firing rate of a cell can act as a readily identifiable marker for that heterogeneity.

Roxin et al. [13] pointed out that the expansive nonlinearity of neuronal input-output relations will turn any normally-distributed variability in the magnitude of synaptic input into a skewed distribution of firing rate outputs, providing a simple explanation for why this distribution should be approximately lognormal (Figure 1B) [10,13]. However, this idea is agnostic to the *source* of input heterogeneity, which could be inherited from external sources, or can be internal to the local population – i.e. due to variations in intrinsic excitability or the degree of local network innervation [14,15]. In fact, there is experimental evidence supporting *both* of these “internal” heterogeneities [16–18]. Cortical pyramidal cells have heterogeneous response profiles to in vivo-like input, both in their excitability and in their sensitivity to fluctuations [19], and highly active neurons in the hippocampus are more likely to develop place fields in a novel environment, which suggests variable excitability during place field assignment in vivo [20]. Higher firing rate pyramidal cells in the cortex are more likely to be connected to each other, forming a “rich club” of high firing rate neurons that receive more input from the local network [21,22]. Rather than independent, we suggest that excitability and local network-related sources of neuronal heterogeneity are related and that this relation arises from, and is maintained by, neuronal dynamics during alternating sleep/wake cycles.

## Biophysical heterogeneity as a point of differential regulation by sleep state

The biophysical properties that set a neuron’s baseline firing rate are precisely those that are influenced by sleep regulatory forces. Sleep-associated loss of neuromodulatory tone changes the channel ecology and extracellular ionic milieu of forebrain neurons [23,24], both of which result in a decrease in neural gain [25,26]. In compensation for the gain-related decrease in excitability, loss of modulatory tone during NREM sleep increases the efficacy of local excitatory synapses and decreases that from excitatory to inhibitory cells [9,26–29], transitioning the population from an externally-driven to an internally-driven regime. Together, these biophysical forces would be expected to narrow the firing rate distribution (Figure 1B), with the relatively large gain-related decrease in spiking of high-firing rate units complemented by a global increase in drive from the local network. High-density electrophysiological recordings confirm that the most pronounced effect of sleep on cortical firing rates is a narrowing of the firing rate distribution [4,30,31]-most prominent during NREM sleep (Figure 1C). It’s important to note that although the shape of the distribution changes, the relative rank of cells in this distribution does not. This distribution narrowing shows a strong initial phase, indicative of the direct state-related effects discussed above, as well as a gradual narrowing over the course of NREM.

In addition to direct neuromodulatory effects, state-specific patterns of population activity can also affect neuronal firing rates. Evidence of such indirect effects is seen in correlations between spiking and the incidence or properties of specific sleep oscillations. Homeostatic changes in firing rate during NREM are correlated with slow wave activity [4,32], indicating that rate homeostasis is related to this prominent activity pattern of the NREM state. The slow oscillation-related decrease in activity of high firing rate neurons persists beyond NREM sleep [4] (Figure 1D), indicating that the slow oscillation imparts a lasting effect on firing rates. These lasting effects are presumably the result of state-specific plasticity [33] – state-specific neuromodulation of synaptic plasticity rules or the interaction of synaptic plasticity rules with state-specific spiking dynamics.

## Homeostatic function of the NREM slow oscillation

By what mechanisms could the NREM slow oscillation enact a lasting narrowing of the neuronal firing rate distribution? Synaptic plasticity is thought to require temporal coordination between synaptically connected neurons [34]. While this may seem straightforward in the stimulus-driven waking state (neurons associated with related stimuli will tend to be co-activated), neuronal activity during sleep is internally generated. This begs the question: how can self-generated activity self-organize to perform a desired plasticity function? Interestingly, spiking during the NREM slow oscillation reveals an intrinsic separation between high and low firing rate units. The slow oscillation is characterized by brief (30-200ms) population-wide DOWN states, after which local neurons begin firing in a population-wide UP state [4,35]. On examination, neurons fire in a statistically reliable sequence at the DOWN->UP transition [36] and a neuron’s place in this ordering is correlated with that neuron’s baseline firing rate, such that neurons with higher firing rates tend to spike before those with lower firing rates [37] (Figure 2). Both biophysical [38] and statistical [37] models indicate that this high-before-low rate neuron effect during transitions from quiescence to active firing states can be attributed to the heterogeneity in input between low and high firing rate neurons [13]. We hypothesize that this sequence also has consequences: spike-timing dependent plasticity (STDP) during the DOWN->UP transition phase of the slow oscillation exhibits differential effects on high-firing vs low-firing neurons. The resulting network modifications can combat an inevitable consequence of synaptic plasticity in a spontaneously active neural population: preferential strengthening of synapses involving cells with higher firing rates.

**Figure 2.**
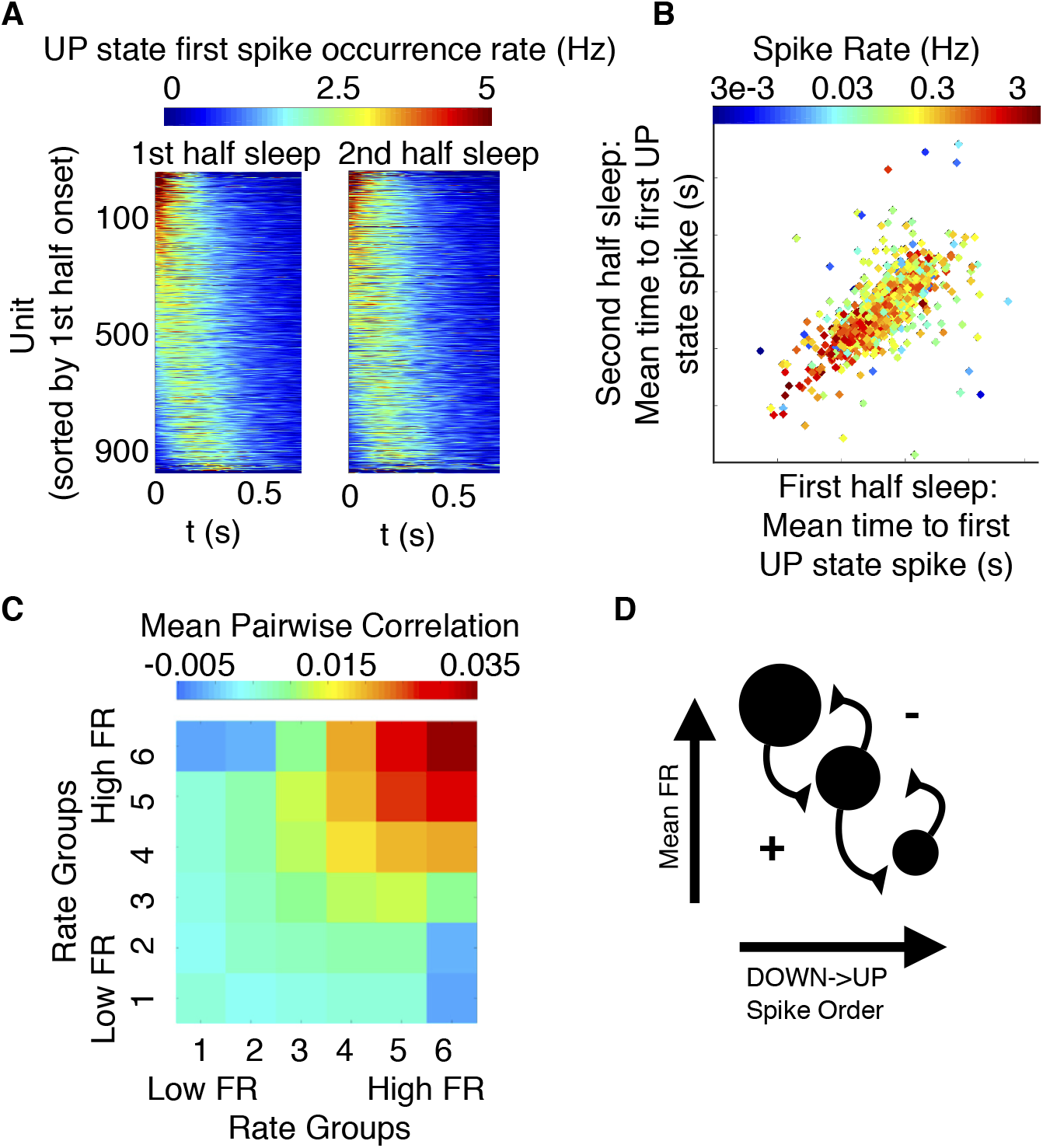
Sequential activity from high to low firing rate cells at the DOWN->UP transition. **A**: Sequential activity at the DOWN->UP transition is consistent across sleep. Peri-event time histogram for the first spike after the UP state onset for first half and second half of sleep from a dataset of cortical excitatory cells. **B**: Onset time is correlated with mean spike rate. **C**: Mean pairwise firing rate correlation between high and low firing rate cells during NREM. **D**: Spontaneous plastic pressure between low (smaller circles) and high (larger circles) firing rate cells during NREM due to high-to-low firing rate sequential activity at the DOWN->UP transition. (Panels A-B from Watson et al 2016)

### Plasticity in a neuronal population with heterogeneous firing rates

Despite the utility of Hebbian plasticity as a neurophysiological mechanism for learning and memory [39], models of neurons connected with Hebbian-plastic synapses demonstrate that networks governed by such a learning rule alone are unstable [40]. This instability is in part because Hebbian plasticity tends to strengthen strong synapses in spontaneously active networks even in the absence of external stimulation-cells that fire together are more likely to fire together again, resulting in a positive feedback loop in which synaptic weights increase ad infinitum [40–42]. While homeostatic plasticity rules can counter the effect of synaptic heterogeneity within a postsynaptic neuron by connecting local (single-synapse) potentiation to global synaptic depression, these rules fail to address a related problem that arises due to *neuronal* heterogeneity.

By definition, neurons obeying Hebbian plasticity rules must spike in order for synapses to be modified, and thus changes in synaptic weight happen at a rate proportional to the number of spikes a neuron fires. The higher the firing rate of the neurons involved, the more chances they have to spike coincidentally, and thus synapses involving high firing rate units have more opportunities for synaptic modification. As a result, spike-timing dependent plasticity (STDP) in an asynchronously active heterogeneous population leads to asymmetric strengthening of synapses to neurons with higher spontaneous firing rates (Box 1). We refer to this drive of spontaneous network dynamics to modify network structure as *spontaneous plastic pressure (SPP)*, in contrast to the more commonly considered “evoked” plastic pressure to change network structure in response to a specific stimulus. From this perspective, SPP is an emergent property of the interaction between synaptic plasticity, neuronal heterogeneity, and population dynamics.

The implication of SPP in an asynchronous (wake-like) population is that variations in neuronal excitability across a population will inevitably lead to a local network structure in which more excitable neurons receive stronger local input than less excitable neurons. While such a heterogeneous network has benefits for neuronal function (see subsequent section), runaway feedback of this effect would destabilize the network due to a widening distribution of neuronal firing rates, and needs to be counterbalanced [15]. The recent experimental work outlined above reveals that firing rate distribution widening is indeed seen during the waking state, and that the NREM slow oscillation is correlated with compensatory distribution narrowing during sleep [4]. We next outline a mechanism by which a single plasticity rule and the experimentally observed spike timing tendencies of the slow oscillation can apply SPP to implement this novel form of homeostasis, which not only accounts for but actually utilizes population heterogeneity to do so.

### Mechanism of homeostatic function of the NREM Slow Oscillation

The sequential pattern of neuronal firing during the DOWN->UP transition provides a putative mechanism by which the NREM slow oscillation can implement such a homeostatic function (Figure 2). Detailed biophysical models indicate that repeated sequential activity at the DOWN->UP transition is strengthened with STDP [43], and this sequential activity is accompanied by increased power in the LFP high gamma (80-100Hz) range [4]-the same time scale needed for synaptic modification via STDP [44,45]. In cortical slice, an LTP protocol during the onset of the UP state was found to effectively facilitate synaptic potentiation following an STDP rule with a minimal number of spike pairings [46]. Because high firing rate neurons tend to fire earlier than low firing rate neurons at the DOWN->UP transition, STDP will tend to increase synaptic weights from high firing rate to low firing rate units, while decreasing weights from low firing rate to high firing rate units (Figure 2D). This redistribution of synaptic weight from high to low firing rate neurons would pull both ends of the firing rate distribution closer to the mean, applying SPP during the synchronous NREM state that acts as a homeostatic counter to that applied in the asynchronous wake state. While sleep-related narrowing of the firing rate distribution combats widening, it does not return the distribution to uniformity, indicating that alternating sleep/wake cycles are a crucial determinant of the general statistics of neuronal network structure and that together the act to maintain a network structure that supports skewed (but not too skewed) population statistics.

### Motivation for homeostatic maintenance of a skewed population

Why maintain a skewed network structure? Theoretical work suggests that skewed populations have beneficial features for network performance. A pool of high firing rate network hubs [21] can facilitate signal transmission between embedded feedforward subnetworks or spike sequences [47,48]. Furthermore, a population of high firing rate “rigid’ and low firing rate “plastic” neurons [6] is beneficial for both system stability and mnemonic function. Stability analysis indicates that network stability relies on a small number of parameter-sensitive high firing rate cells, allowing the properties of many “sloppy” low firing rate cells to each change a great deal with learning without a negative impact on network dynamics [49]. These plastic network elements alone would allow for strong memories that are quickly formed but quickly overwritten, while rigid elements alone result in long-lasting but weak memories [50]. Together, a distributed memory system with the ability to transfer memory information between rigid and plastic elements maximizes both capacity and stability [51]. Memory consolidation during the NREM slow oscillation provides an opportune window for such rigid-plastic interaction.

## Consolidation: the interaction of local and global plasticity during the NREM slow oscillation

According to the “Two Stage Model” [52], consolidation involves the transfer of memory traces from short-term storage in the hippocampus to long term storage in the neocortex [53,54]. Transfer is thought to occur during hippocampal Sharp-Wave Ripple events (SPW-R)[55]. SPW-Rs are brief (∼100ms) population bursts in hippocampal CA1 that are associated with “replay” of task-related patterns of spiking activity in both the hippocampus and neocortex and correlate with memory task performance [56–58]. Recent evidence suggests that the NREM slow oscillation is an integral part of the cortical response to SPW-Rs. SPW-Rs are temporally coordinated with slow wave activity [59], and have been found to both precede [58,60] and follow [61] the cortical DOWN state. In a closed-loop experiment using SPW-R—triggered electrical stimulation of slow waves in the neocortex, induced ripple-slow wave events were found to improve memory performance of novel objects [62]. During a subset of these evoked events, the “usual” sequential activation at the DOWN->UP transition was altered. Importantly, neurons that spiked at a different time in the sequence developed recognition fields to the novel object. This suggests that these neurons with perturbed activity were the beneficiaries of specific memory-related plasticity.

In order for hippocampal replay to transfer a memory trace to a cortical network, the neocortical population response must result in the integration of a novel pattern into the existing local network structure. Replay-induced perturbation of the sequential activation from high to low firing rate neurons at the DOWN->UP transition presents a window of opportunity for this integration, possibly by strengthening synapses from rigid high firing rate cells to specific memory-related, low firing rate plastic neurons tagged by hippocampal replay events (Figure 3). In this way, the coupling of mnemonic and homeostatic function at the DOWN->UP transition could act as a vehicle for the interaction between local and global plasticity.

**Figure 3.**
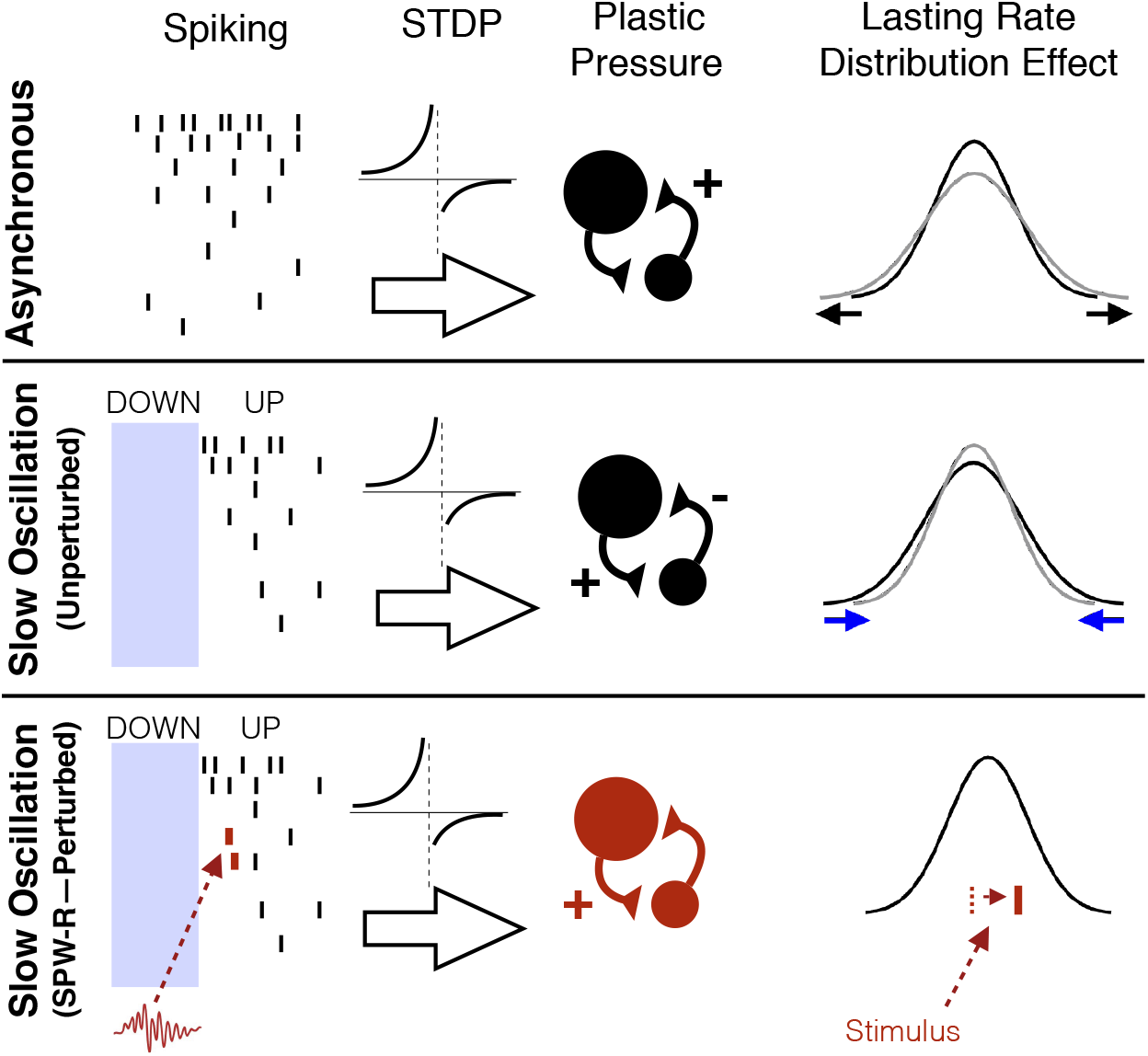
Summary: Spontaneous plastic pressure in different contexts. STDP during asynchronous spiking applies SPP that strengthens synapses to high firing rate units, widening the firing rate distribution. STDP during sequential high-to-low rate spiking during the NREM DOWN->UP transition applies SPP that tends to strengthen synapses from high firing rate units to low firing rate units. STDP during SPW-R-mediated perturbation of sequential activity at the DOWN->UP transition strengthens memory-specific synapses, resulting in stimulus-specific spiking at future stimulus presentations.

## Conclusions

The self-generated organization of neuronal activity during sleep promotes both network stabilization and mnemonic functions, which are achieved through the interaction between synaptic plasticity and state-specific patterns of population dynamics. Recent findings suggest a framework in which neuronal dynamics during the NREM slow oscillation act as a locus of both homeostatic and mnemonic functions. By this model, both functions are the result of synaptic plasticity during sequential activation from higher to lower firing rate neurons at the DOWN->UP transition. We hypothesize that the unperturbed DOWN->UP transition enacts homeostatic maintenance of the neuronal firing rate distribution. This novel form of homeostasis would compensate for the widening of the firing rate distribution due to spontaneous plastic pressure applied by asynchronous activity during wakefulness. Perturbation of spike timing at the DOWN->UP transition by hippocampal ripples, on the other hand, enables hippocampus-mediated memory consolidation, serving to integrate novel memory traces into the existing cortical framework while allowing for a more general homeostatic effect to occur in the background.

### Box 1

Plastic pressure in an asynchronous state

How may we describe mathematically the plastic pressure applied by spontaneous activity in an asynchronous (wake-like) population of neurons with heterogeneous firing rates? Here we present a basic calculation of the spontaneous plastic pressure (SPP) on a synapse in such a population, and investigate the asymmetry (i.e. the difference in SPP applied to a synapse from a low to a high rate cell compared to one in the opposite direction) due to three relevant variables: 1) spontaneous firing rates of the pre and postsynaptic cells, 2) current synaptic weights and 3) relative strength of potentiation/depression in the synaptic plasticity rule. We show that, while SPP does not itself create asymmetry in synapses between high and low firing rate neurons, it does accentuate asymmetries that exist, both in the synaptic weights and in the plasticity rule, in a way that favors synapses to and between neurons with higher firing rate.

We first define plastic pressure on synapse *ij* from presynaptic neuron *j* to postsynaptic neuron *i* as the expected rate of change of synaptic weight,

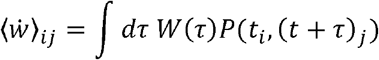

where *P*(*t_i_*,(*t* + τ)_*j*_) = *P*(*t*_*i*_) *P*((*t* + τ)_*j*_|*t_i_*) is the probability of neurons *i* and *j* co-spiking with small time delay τ and *W*(τ) is a plasticity rule that gives the change in synaptic weight as a function of delay τ between spike times [63]. Suppose that the dominant sources of rate correlation are synaptic interactions (i.e. the population is in an “asynchronous” state [64]). We represent their effect by adding *K_ij_*(τ) to the cross-correlation, i.e. *P*((*t* + τ)_*j*_|*t_i_*) = *P*(*t*_*j*_) + *K_ij_*(τ) assuming that *K_ij_*(τ) is small. The expression for plastic pressure then simplifies to

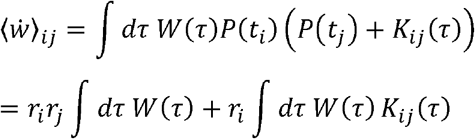

in which the first term, SPP due to asymmetry in the STDP rule, depends on the rates of both neuron *i* and neuron *j*, while the second term, SPP from synaptic factors, depends on the *rate of the postsynaptic neuron i only*. There are two contributions to the synaptic correlation term, *K_ij_*(τ). One, *w_ij_k*(τ), for spikes added to neuron *i* due to synapse *ij*, and one for spikes added to neuron *j* due to synapse *ij*, *w_ij_k*(–τ), giving us: *K_ij_*(τ) = *w_ij_k*(τ) + *w_ij_k*(–τ). With this recognition we express the pressure on synapse *ij* as

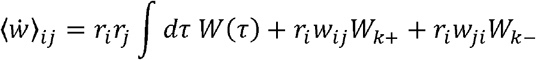

where *W*_*k*+_ = ∫ *d*τ *W*(τ)*k*(τ) and *W*_*k*−_ = ∫ *d*τ *W*(τ)*k*(–τ) are the plastic pressure applied by a unitary synapse in each direction due to the plasticity rule. The term with *W*_*k*+_ represents the positive SPP on synapse *ij* due to synapse *ij*; it is proportional to the firing rate of neuron *i* and the current weight *w_ij_*. The term with *W*_*k*−_ corresponds to the negative SPP on synapse *ij* due to synapse *ji* in the reverse direction, and is proportional to the firing rate of neuron *i* and the current weight *w_ji_*.

We now ask, *how much more will SPP during asynchronous activity strengthen synapses from lower firing rate neurons to higher firing rate neurons than those in the reverse direction?* For this, we calculate the asymmetry in pressure applied to the synapse *ij* compared to that applied to *ji*, as a function of the firing rates, weights, and the unitary plastic pressure:

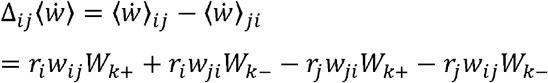

We illustrate schematically below the relative SPP between two neurons of different firing rates for 3 simplified cases. In **Case 1**, with equal synaptic weights and a symmetric learning rule (i.e. *w_ij_* = *w_ij_* and *W*_*k*+_=-*W*_*k*-_), plastic pressure is symmetric (i.e. 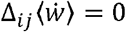). in **Case 2**, with unequal synaptic weights (*w_ij_* ≠ *w_ji_*), plastic pressure will strengthen the higher weight synapse more than the lower weight synapse, and this effect will be stronger for higher firing rate pairs. Finally, in **Case 3** with an asymmetric learning rule (i.e. W_k+_ >-W_k-_, as is observed experimentally [65]), SPP will asymmetrically strengthen synapses onto higher firing rate neurons. Together, these results indicate that synapses *to* and *between* high firing rate neurons will be preferentially strengthened by asynchronous spontaneous activity.

**Box figure.**
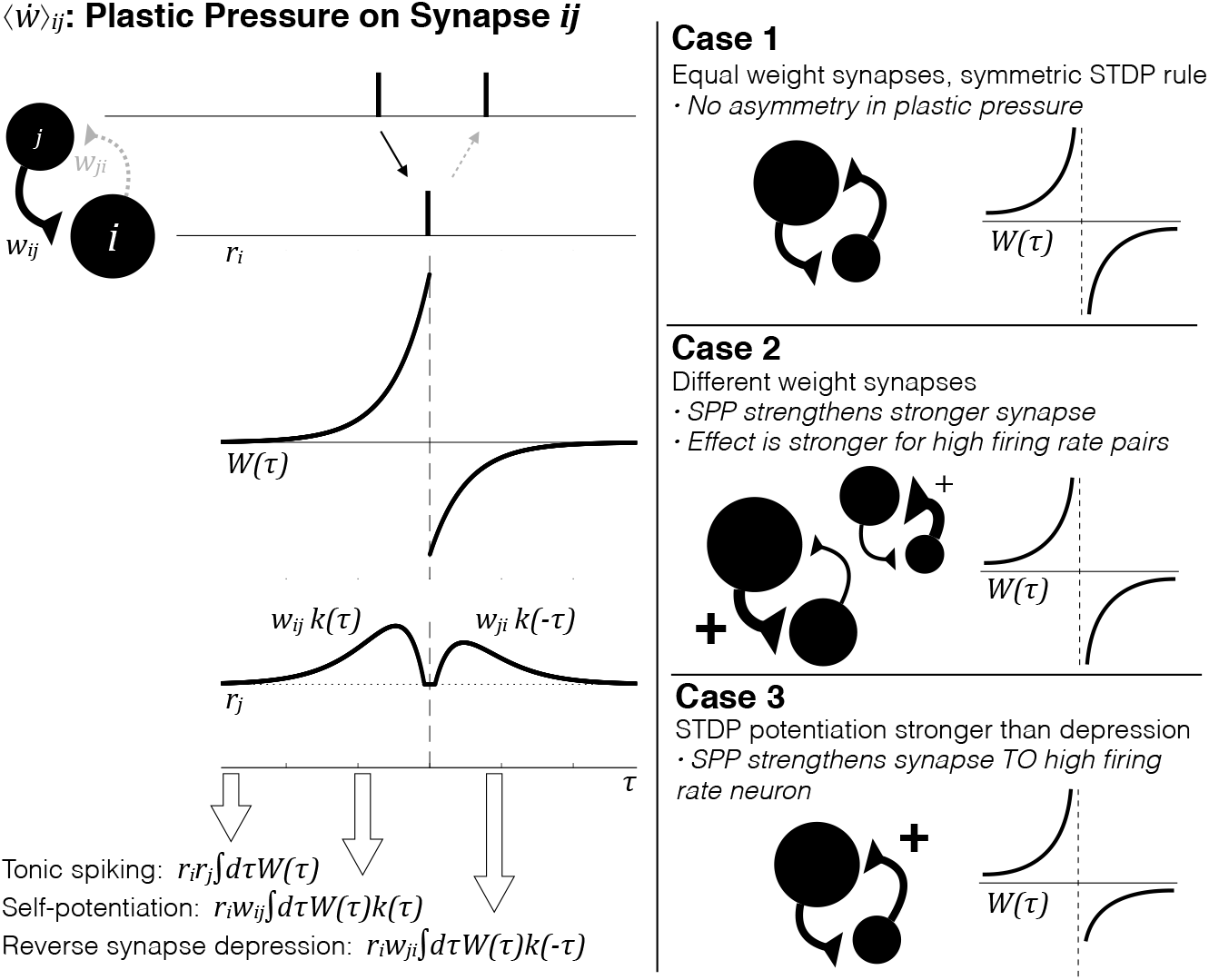
Spontaneous plastic pressure (SPP) due to asynchronous activity. (Left) Calculation of SPP on synapse from presynaptic neuron j to postsynaptic neuron i. (Right) Schematic of the SPP asymmetry 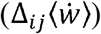 between a representative bidirectionally-connected high firing rate (large circles) and a low firing rate (smaller circles) neuron pair for three illustrative cases, as a function of the synaptic weights, STDP rule, and relative firing rate.

## Acknowledgements

The authors would like to thank Rachel Swanson for extensive comments during writing of the manuscript, Andres Grosmark for comments and ongoing discussion of relevant topics, and all members of the Buzsáki lab for continuously insightful discussion.

